# Statin use and breast cancer survival: A Swedish nationwide study

**DOI:** 10.1101/335034

**Authors:** Signe Borgquist, Per Broberg, Jasaman Tojjar, Håkan Olsson

## Abstract

**Background:** A sizeable body of evidence suggests that statins can cease breast cancer progression and prevent breast cancer recurrence. The latest studies have, however, not been supportive of such clinically beneficial effects. These discrepancies may be explained by insufficient power. This considerably sized study investigates the association between both pre- and post-diagnostic statin use and breast cancer outcome.

**Methods:** A Swedish nation-wide retrospective cohort study of 20,559 Swedish women diagnosed with breast cancer (July 1^st^, 2005 through 2008). Dispensed statin medication was identified through the Swedish Prescription Registry. Breast cancer related death information was obtained from the national cause-of-death registry until December 31^st^, 2012. Cox regression models yielded hazard ratios (HR) and 95% confidence intervals (CI) regarding associations between statin use and breast cancer-specific and overall mortality.

**Results:** During follow-up, a total of 4,678 patients died, of which 2,669 were considered breast cancer related deaths. Compared to non- or irregular use, regular pre-diagnostic statin use was associated with lower risk of breast cancer related deaths (HR=0.77; 95% CI 0.63–0.95, P=0.014). Similarly, post-diagnostic statin use compared to non-use was associated with lower risk of breast cancer related deaths (HR=0.83; 95% CI 0.75–0.93, P=0.001).

**Conclusion:** This study evidently supports the notion that statin use is protective regarding breast cancer related mortality in agreement with previous Scandinavian studies, although less so with studies in other populations. These disparities should be further investigated to pave the way for future clinical trials investigating the role of statins in breast cancer.

## Introduction

Breast cancer tends to occur in older women with a median age at diagnosis of 60 years in developed countries. Consequently, many breast cancer patients may be treated for both pre- and post-diagnostic comorbidities. Frequently prescribed drugs are statins, which are cholesterol-lowering HMG-CoA reductase inhibitors used for prevention of cardiovascular diseases. In Sweden, around 20% of the adult female population is currently prescribed a statin according to The National Board of Health and Welfare of Sweden (www.socialstyrelsen.se). Statins block the rate-limiting step in the mevalonate pathway by inhibiting HMG-CoA reductase, predominantly in hepatocytes ^1^. The reduction in intracellular cholesterol levels in the liver induces up-regulation of the low-density-lipoprotein receptor resulting in lower levels of circulating cholesterol. Beyond cholesterol metabolism, the mevalonate pathway is essential for tumor promoting effects of the oncogene p53 ^2^. Additionally, isoprenoid production plays a fundamental role in cell growth regulation^3^. Recent observational studies have reported an inverse association between pre-diagnostic statin use and risk of breast cancer-related death ^4, 5^. These findings have, however, been replicated to a lesser degree in studies performed outside of Scandinavia ^6, 7^ A recent systematic review summarized relative risks of statin use with breast cancer recurrence ^8^. Given the epidemiological findings from other Scandinavian populations, we hypothesized that statins may also have anticancer effects and therefore reduce cancer related mortality in a Swedish population. In this nationwide study, we investigated breast cancer related and all-cause mortality among breast cancer patients according to statin use prior to and/or following their breast cancer diagnosis.

## Material and Methods

### Study population

All breast cancer cases were identified through the Swedish Cancer Registry, which has a high level of completeness and is considered to be of good quality as approximately 99% of the cases have been morphologically verified. Incident breast cancer cases were identified in the registry using the code ICD7=170* (according to the International Classification of Diseases). Men were excluded from this study given the low incidence of male breast cancer. Breast cancer patients <40 years of age at diagnosis were excluded from the analyses since this age group is unlikely to receive statins, and if so, were considered as being in a certain high-risk group (such as familial hypercholesterolemia). Moreover, women were excluded if any previous cancer diagnosis was recorded from the initiation of the Swedish Cancer Registry in 1958 until the end of year 1999. Thus, the study included 20,559 women with breast cancer diagnosed in Sweden from July 1^st^, 2005 through 2008 (Table 1). Dates for birth, breast cancer diagnosis, and TNM-stages were extracted from the cancer registry along with the actual cases. The Swedish Cancer Registry started recording TNM information in 2004 although there were some missing data at the beginning of its launching period. Pharmaceutical records, allowing for information on parameters such as statin prescriptions, are available from July 1^st^, 2005 when the Swedish Prescribed Drug Registry initiated nationwide retrieval of all prescribed drugs. In this study, when evaluating pre-diagnostic statin use, we investigated breast cancer cases diagnosed thereafter with a minimum of six months delay (that is January 1^st^, 2006.). In the analysis of regular statin usage, the data set was further restricted to patients diagnosed with breast cancer after 1^st^ July 2007.

**Table 1:**
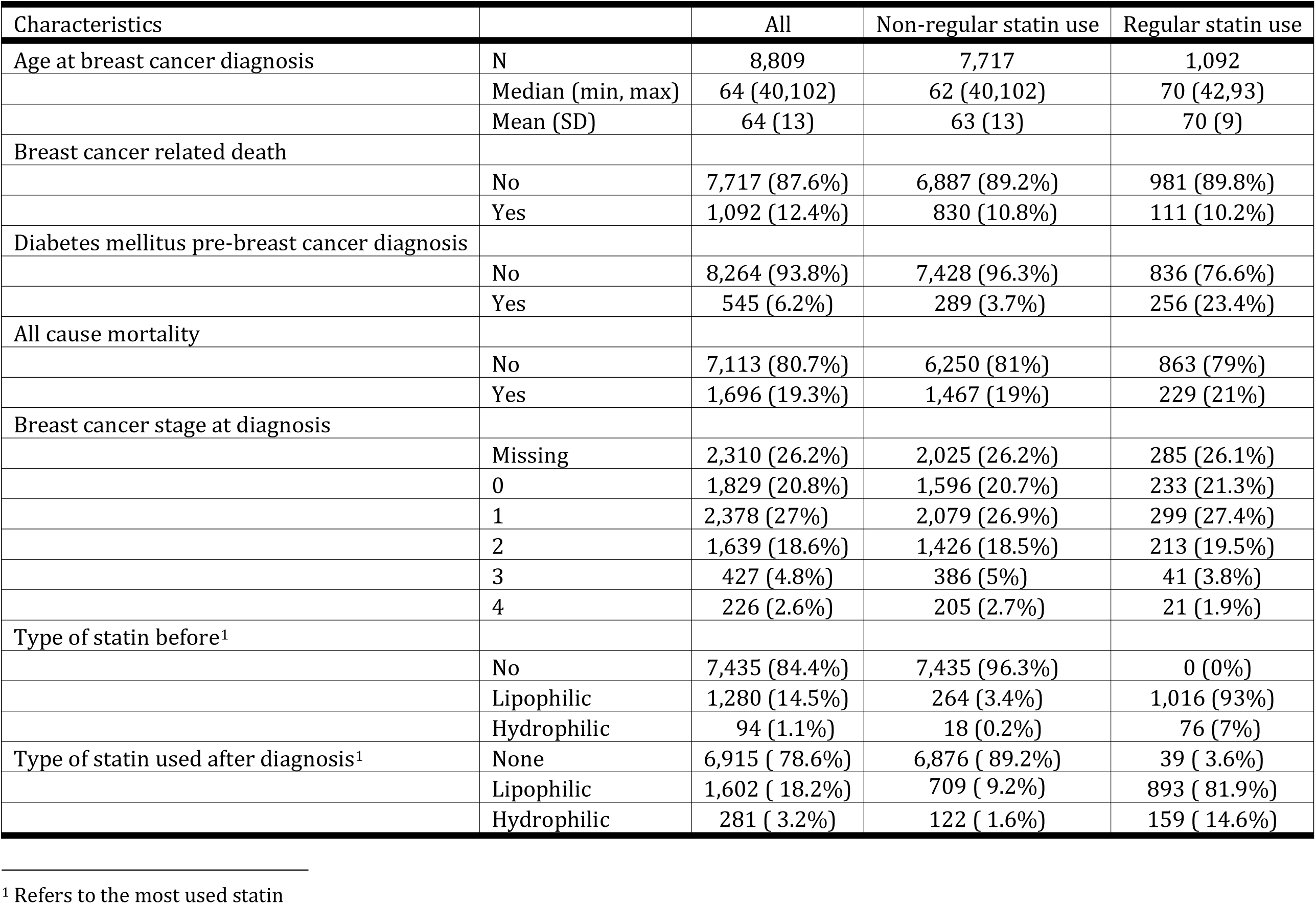
Patient – and disease characteristics according to regular statin use among Swedish women diagnosed with breast cancer at age 40+ from July 1^st^, 2007 – December 31^st^, 2008. They were followed up until December 31^st^, 2012 for breast cancer related death, and, December 31^st^, 2013, for all-cause mortality.

| Characteristics |  | All | Non-regular statin use | Regular statin use |
| --- | --- | --- | --- | --- |
| Age at breast cancer diagnosis | N | 8,809 | 7,717 | 1,092 |
|  | Median (min, max) | 64 (40,102) | 62 (40,102) | 70 (42,93) |
|  | Mean (SD) | 64 (13) | 63 (13) | 70 (9) |
| Breast cancer related death |  |  |  |  |
|  | No | 7,717 (87.6%) | 6,887 (89.2%) | 981 (89.8%) |
|  | Yes | 1,092 (12.4%) | 830 (10.8%) | 111 (10.2%) |
| Diabetes mellitus pre-breast cancer diagnosis |  |  |  |  |
|  | No | 8,264 (93.8%) | 7,428 (96.3%) | 836 (76.6%) |
|  | Yes | 545 (6.2%) | 289 (3.7%) | 256 (23.4%) |
| All cause mortality |  |  |  |  |
|  | No | 7,113 (80.7%) | 6,250 (81%) | 863 (79%) |
|  | Yes | 1,696 (19.3%) | 1,467 (19%) | 229 (21%) |
| Breast cancer stage at diagnosis |  |  |  |  |
|  | Missing | 2,310 (26.2%) | 2,025 (26.2%) | 285 (26.1%) |
|  | 0 | 1,829 (20.8%) | 1,596 (20.7%) | 233 (21.3%) |
|  | 1 | 2,378 (27%) | 2,079 (26.9%) | 299 (27.4%) |
|  | 2 | 1,639 (18.6%) | 1,426 (18.5%) | 213 (19.5%) |
|  | 3 | 427 (4.8%) | 386 (5%) | 41 (3.8%) |
|  | 4 | 226 (2.6%) | 205 (2.7%) | 21 (1.9%) |
| Type of statin before<sup>1</sup> |  |  |  |  |
|  | No | 7,435 (84.4%) | 7,435 (96.3%) | 0 (0%) |
|  | Lipophilic | 1,280 (14.5%) | 264 (3.4%) | 1,016 (93%) |
|  | Hydrophilic | 94 (1.1%) | 18 (0.2%) | 76 (7%) |
| Type of statin used after diagnosis<sup>1</sup> | None | 6,915 ( 78.6%) | 6,876 ( 89.2%) | 39 ( 3.6%) |
|  | Lipophilic | 1,602 ( 18.2%) | 709 ( 9.2%) | 893 ( 81.9%) |
|  | Hydrophilic | 281 ( 3.2%) | 122 ( 1.6%) | 159 ( 14.6%) |

<sup>1</sup> Refers to the most used statin

Information on TNM-stage was retrieved from the Swedish Cancer Registry summarizing breast cancer into stages 0–IV according to the international staging system. To account for missing information regarding stage, a separate missing factor level was introduced. Information regarding the cause and date of death was retrieved from the Swedish Cause of Death Register and the Swedish Population Register. Follow-up data up until the 31^st^ of December 2012, respectively 31^st^ of December 2013, were retrieved. Due to lack of the cause of death from 2013, the follow-up time after the 31^st^ of December 2012 was censored. Censoring would occur in case of death due to any cause other than breast cancer prior to this date. Ethical permission was obtained from The Regional Ethical Review Board in Lund, Sweden (Dnr 2013/787).

### Statin use

Statin use was obtained from the Swedish Prescription Registry. Patients were classified as exposed to statins if they had at least one statin prescription logged in their pharmaceutical record with an Anatomic Therapeutic Chemical code (ATC code) beginning with “C10AA”. Consequently, women prescribed a statin at least once were classified as statin-users, whereas others were assumed to be unexposed to statins and therefore classified as non-users. Three types of statin-users were defined: 1) regular, pre-diagnostic statin users, 2) irregular, pre-diagnostic statin users, and 3) post-diagnostic statin users. The regular pre-diagnostic users were defined as women with a statin prescription within six months before their breast cancer diagnosis along with another prescription during the two years prior to the diagnosis. The irregular statin users were defined as statin users not fulfilling the regular-user criteria stated above. Post-diagnostic statin users included all women with any statin prescription following their breast cancer diagnosis irrespective of pre-diagnostic statin use. By using each patient’s unique Civil Personal Registry number, the National Board of Health and Welfare merged retrieved data from the four registries containing the individual’s information (the Swedish Cancer Registry, the Swedish Prescribed Drug Registry, the Swedish Population Register, and the Swedish Cause of Death Registry). For confidentiality reasons, the selected data were then anonymized and returned with a new serial number instead of the civil register number. The identifier for the patient codes was kept at the National Board of Health and Welfare.

### Statistical analyses

The main purpose was to compare mortality, both breast cancer related and overall deaths, between statin and non-statin users. Additionally, the impact of statin dose was assessed. To make different statin doses comparable across the various statins, the World Health Organization-recommended daily dose definition was applied ^9^, and a defined daily dose (DDD) was calculated for all statin users. Based on the penultimate statin prescription prior to the breast cancer diagnosis and the time until the ultimate prescription (which will be the expected time of usage of the penultimate dose), a rate of usage was defined by dividing the dose by the time calculated in days. Thus, comparisons of usage could be made in terms of the amount of daily doses, which were classified into three categories: 1.) 0?0.1; 2.) >0.01?0.5; and 3.) >0.5. In the analysis of post diagnostic usage we used a model that assumes that the effect is related to the logarithm of the cumulative dose such that an increase in that logarithm translates into an exponent affecting the hazard ratio. An increase in cumulative dose from 20 to 1000 DDD will raise the hazard ratio to the power of log(1001)- log(21) = 3.86, assuming everything else being equal. So a hazard ratio of 0.85 becomes 0.85**(log(1001)-log(21)) = 0.53. This model was chosen based on Gasparrini *et al*. ^10^, where data suggest a log-transformation of exposure.

Two types of Cox regression were done: 1.) A standard analysis with baseline characteristics at inception was applied for analyses evaluating associations between pre-diagnostic statin use and clinical outcome and 2.) A Cox regression model allowing for time and varying covariates was used in the analysis of post diagnostic use. The latter approach made it possible for a subject to provide data both as a non- and as statin user. All multivariable standard Cox regression analyses (time dependent excluded) were adjusted for age at diagnosis, breast cancer stage (0?IV), and diabetes diagnosed prior to breast cancer diagnosis. Pre-diagnostic statin usage and mortality were analyzed using two different approaches: 1.) Evaluating the impact of regular usage compared to non-usage and/or 2.) Evaluating the effects of usage rate. In this method, survival time was measured in years since diagnosis. In the second type of analysis, a Cox proportional hazards model with time varying covariates (Cox regression with late entry) was used to allow for medication changes along the way and to assess the impact of cumulative doses. In this case, the time scale was age. In the analysis of the effects of cumulative number of statin prescriptions, a Cox regression with time and varying covariates was employed after log transformation of the number of prescriptions plus one. A sensitivity analysis addressed exposure lag-time to control for potential protopathic bias ^11^. One year was decided as lag-time based on our calculations lagging exposure, or rather set exposure less than a certain amount equal to zero, not with respect to time, but with respect to cumulative dose. The results suggest that the effect rises to a plateau at one year (Supplemental Figure 1). We also calculated the effect of a varying lag-time (Supplemental Figure 2). However, the follow-up in our study cannot exceed eight years, so that defines the range. As may be expected the hazard ratio decreased as the lag increased. Using no lag will likely yield a conservative estimate according to this analysis. This agrees with the intuition, since time periods where the drug may not yet have had an effect still counts in the calculation, when no lag is applied.

**Figure 1:**
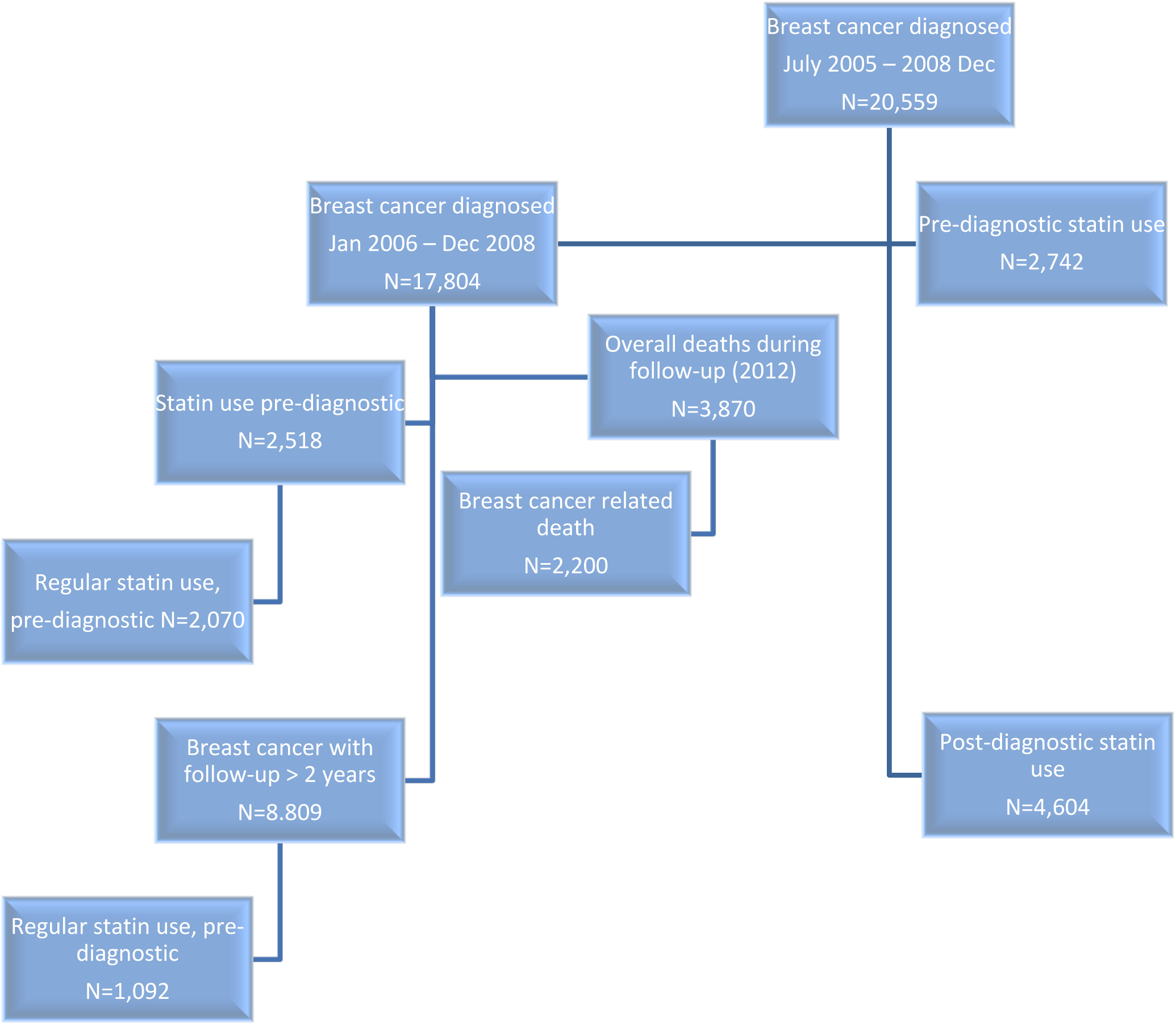
Flow chart illustrating the study population

All statistical analyses were performed using R version 3.2.1 (R Core Team, 2015). Two-tailed P-values were used.

## Results

This nationwide study included 20,559 Swedish women diagnosed with breast cancer after 40 years of age between July 1^st^, 2005 and through 2008, Fig. 1. The average age at diagnosis was 64 years. During follow-up until December 31^st^, 2012, a total of 2,669 breast cancer-related deaths occurred among the 20,559 women, who were diagnosed after 1^st^ January 2006, (Table 2). In this entire study cohort, 2,518 patients were prescribed a statin before their breast cancer diagnosis. For analyses of regular use and prognosis, the cohort of women with more than two years of follow-up were used (N=8,809), and among these 1,092 were regular statin users (Table 1). Simvastatin was the most frequently prescribed statin accounting for 82% (among the 17,804 patients considered for pre-diagnostic statin use) of all recorded statin prescriptions followed by pravastatin, atorvastatin, rosuvastatin, and fluvastatin. The numbers for post-diagnostic use similarly showed that simvastatin was the most frequently prescribed statin (93%) followed by pravastatin, rosuvastatin, lovastatin, and atorvastatin. When comparing use of pre- and post-diagnostic statins, the majority of pre-diagnostic statin users would expectedly also be post-diagnostic statin users (N=2,503/2,739, 91.3%). In addition, a substantial numbers of the non-pre-diagnostic statin users would become users of statins in the post-diagnostic setting (N=2,082/17,801, 11.7%).

**Table 2:** Patient- and disease characteristics according to breast cancer death among Swedish women diagnosed with breast cancer at age 40+ from July 1^st^, 2005 to December 31^st^, 2008. They were followed up until December 31^st^, 2012 for breast cancer related death, and, December 31^st^, 2013 for all-cause mortality.

| Characteristics |  | All | Not dead in breast cancer | Breast cancer related death |
| --- | --- | --- | --- | --- |
| Age at breast cancer diagnosis | N | 20,559 | 17,890 | 2,669 |
|  | Median (min, max) | 63 (40,102) | 62 (40,101) | 75 (40,102) |
|  | Mean (SD) | 64 (13) | 63 (12) | 72 (14) |
| Regular statin user |  |  |  |  |
|  | No | 7,717 (87.6%) | 6,887 (87.5%) | 830 (88.2%) |
|  | Yes | 1,092 (12.4%) | 981 (12.5%) | 111 (11.8%) |
| Diabetes mellitus pre-breast cancer diagnosis |  |  |  |  |
|  | No | 19,343 (94.1%) | 16,957 (94.8%) | 2,386 (89.4%) |
|  | Yes | 1,216 (5.9%) | 933 (5.2%) | 283 (10.6%) |
| All cause mortality |  |  |  |  |
|  | No | 15,881 (77.2%) | 15,881 (88.8%) | 0 (0%) |
|  | Yes | 4,678 (22.8%) | 2,009 (11.2%) | 2,669 (100%) |
| Breast cancer stage at diagnosis |  |  |  |  |
|  | Missing | 7,383 (35.9%) | 6,247 (34.9%) | 1,136 (42.6%) |
|  | 0 | 3,506 (17.1%) | 3,394 (19%) | 112 (4.2%) |
|  | 1 | 4,818 (23.4%) | 4,640 (25.9%) | 178 (6.7%) |
|  | 2 | 3,419 (16.6%) | 2,882 (16.1%) | 537 (20.1%) |
|  | 3 | 855 (4.2%) | 556 (3.1%) | 299 (11.2%) |
|  | 4 | 578 (2.8%) | 171 (1%) | 407 (15.2%) |
| Type of statin before breast cancer diagnosis |  |  |  |  |
|  | No | 17,817 (86.7%) | 15,507 (86.7%) | 2,310 (86.5%) |
|  | Lipophilic | 2,549 (12.4%) | 2,210 (12.4%) | 339 (12.7%) |
|  | Hydrophilic | 193 (0.9%) | 173 (1%) | 20 (0.7%) |

### Statin use and breast cancer related mortality

#### Regular pre-diagnostic statin use

Compared to non- or irregular statin use, regular statin use was associated with lower risk of breast cancer related deaths (HR=0.77; 95%CI 0.63?0.94, P=0.014), stratified for age at diagnosis and tumor stage, adjusted for diabetes (Table 3). When evaluating only diabetic patients (N=545), the association was strengthened (HR=0.63; 95%CI 0.40?0.99, P=0.044). The breast cancer survival effects of different statin doses were evaluated using the calculated daily statin dose. With the lowest daily dose (0?0.1) serving as reference, use of higher statin daily doses (>0.1?0.5) was associated with fewer breast cancer related deaths (HR=0.74; 95%CI 0.58?0.95, P=0.019). The highest daily dose (>0.5) also improved breast cancer related survival significantly compared to the lowest dose (HR=0.84; 95%CI 0.72?0.98, P=0.022), Table 3.

**Table 3:** Breast cancer related death and overall death in relation to statin use prior to breast cancer diagnosis^1^.

|  | Breast cancer related death |  |  |  |  | Overall death |  |  |  |  |
| --- | --- | --- | --- | --- | --- | --- | --- | --- | --- | --- |
| Pre-diagnostic statin use | Events | Person years | HR | 95% Confidence Intervals | P-value | Events | Person years | HR | 95% Confidence Intervals | P-value |
| **Statin use rate** |  |  |  |  |  |  |  |  |  |  |
| None (n=15,718) | 1,936 | 77,604 | 1.0 | - | - | 3,358 | 89,972 | 1.0 |  |  |
| Intermed. (n=576) | 65 | 2,786 | 0.74 | 0.58 - 0.95 | 0.019 | 297 | 3,229 | 0.75 | 0.63 - 0.89 | 0.001 |
| High (n=1,492) | 189 | 7,191 | 0.84 | 0.72 - 0.98 | 0.022 | 216 | 8,303 | 0.80 | 0.72 - 0.89 | <0.001 |
| **Regular statin use** |  |  |  |  |  |  |  |  |  |  |
| No (n=7,717) | 830 | 33,321 | 1 | - | - | 1,467 | 39,566 | 1 | - | - |
| Yes (n=1,092) | 111 | 4,703 | 0.77 | 0.63 - 0.95 | 0.014 | 229 | 5,566 | 0.76 | 0.66 - 0.88 | <0.001 |

<sup>1</sup> The Cox regression took age at diagnosis, stage and diabetes into account.

#### Post-diagnostic statin use

Statin use following breast cancer diagnosis was associated with lower risks of breast cancer related deaths compared to non-users (HR=0.83; 95%CI 0.75?0.93, P=0.001), as shown in Table 4. A sensitivity analysis considering one-year lag time showed a similar association (HR=0.94; 95%CI 0.92?0.96, P<0.001).

**Table 4:** Breast cancer related death and overall death in relation to statin use following breast cancer diagnosis1.

|  | Breast cancer related death |  |  |  |  | Overall death |  |  |  |  |
| --- | --- | --- | --- | --- | --- | --- | --- | --- | --- | --- |
| Post-diagnostic statin use | Events | Person years | HR | 95% Confidence Intervals | P-value | Events | Person years | HR | 95% Confidence Intervals | P-value |
| **Any statin use** |  |  |  |  |  |  |  |  |  |  |
| No (n=20,503) | 2,245 | 86856 | - | - |  | 3,811 | 99102 |  |  |  |
| Yes (n=4,358) | 412 | 17715 | 0.83 | 0.75 – 0.93 | 0.001 | 848 | 21340 | 0.89 | 0.82 – 0.96 | 0.003 |
| **Daily statin dose** |  |  |  |  |  |  |  |  |  |  |
| - |  |  |  |  |  |  |  |  |  |  |
| log (1+ddd) (n=93,595) | 2,657 | 104571<sup>2</sup> | 0.93 | 0.91 – 0.95 | <0.001 | 4,659 | 120442 | 0.96 | 0.95 – 0.97 | <0.001 |
| **Daily statin dose by solubility** |  |  |  |  |  |  |  |  |  |  |
| Lipophilic log(1+ddd) |  |  | 0.93 | 0.91 – 0.96 | <0.001 |  |  | 0.96 | 0.95 – 0.97 | <0.001 |
| Hydrophilic log(1+ddd) |  |  | 0.93 | 0.89 – 0.98 | 0.002 |  |  | 0.96 | 0.93 – 0.99 | 0.003 |
| Other log(1+ddd) |  |  | 0.99 | 0.84 – 1.17 | 0.93 |  |  | 1.07 | 0.97 – 1.16 | 0.24 |

<sup>1</sup> The Cox regression age at diagnosis, stage and diabetes took into account.
<sup>2</sup> In the analysis of defined daily statin dose, the number of person years has been calculated for all subjects included without regard to treatment status. The same goes for the number of events.

### Statin use and all cause of mortality

#### Pre-diagnostic statin use

All-cause mortality according to pre-diagnostic statin use provided results similar to breast cancer related mortality with a lower risk of death among regular statin users compared to non-regular users (HR=0.76; 95%CI 0.66?0.88, P<0.001) as shown in Table 3.

The overall survival effects of different statin doses were evaluated using the calculated statin use rate. With no statin use serving as the reference, the intermediate statin use rate was associated with fewer deaths (HR=0.75; 95%CI 0.63?0.89), P=0.001). Accordingly, the highest dose decreased all-cause mortality significantly compared to the lowest dose (HR=0.80; 95%CI 0.72–0.89, P<0.001), shown in Table 3.

#### Post-diagnostic statin use

All-cause mortality among the post-diagnostic statin user was similar to breast cancer related mortality with a lower risk of overall death among statin users compared to non-regular statin users (HR=0.89; 95% CI 0.82?0.96, P=0.003) as shown in Table 4. Similarly, daily statin use was associated with improved overall survival (HR=0.96; 95% CI 0.95?0.97, P<0.001). These associations held true for both lipophilic and hydrophilic statin use as shown in Table4. A sensitivity analysis considering one-year lag time strengthened the association (HR=0.95; 95%CI 0.93?0.96, P<0.001).

## Discussion

This large-scaled nationwide Swedish study demonstrates that the use of cholesterol-lowering statins is associated with lower risk of breast cancer related and overall deaths among women diagnosed with breast cancer, irrespective of whether statins were used pre- or post-diagnosis. These results confirm previously presented studies based on Scandinavian cohorts ^4, 5, 9^ although the results have been less evident in studies from England ^12^, Ireland ^7^, and Scotland ^6^. In line with our results, a recent comprehensive meta-analysis addressing statin use and breast cancer survival demonstrated significant survival benefits among statin users, both in terms of overall survival and disease-specific survival, which was true for both pre- and post-diagnosis statin use ^13^. Another systematic review on statin use and cancer mortality, including breast cancer mortality, showed consistent findings ^14^. In summary, consistent observational evidence support a reduced risk of breast cancer recurrence/mortality among statin users ^4, 5, 15-18^, which have paved the way for multiple ongoing clinical trials, which are investigating the role of statins in breast cancer (such as #NCT02483871, #NCT02958852, #NCT01988571). Nevertheless, treatment predictive biomarkers enabling a more targeted approach and selection of patients likely to respond to statin treatment is required ^19^. Recently, a novel multigene signature comprised of genes involved in the cholesterol biosynthesis, was shown to predict statin sensitivity ^20^.

Our current findings are biologically plausible since statins inhibit not only cholesterol synthesis but also reduce other important downstream products, of which several are used in cell proliferation such as membrane integrity maintenance, cell signaling, protein synthesis, and cell-cycle progression ^21-23^. Interruption of these processes in potentially remnant malignant cells following primary breast cancer may therefore inhibit further cancer growth and metastasis as depicted by an improved clinical outcome. Not only can statins have a direct impact on cancer cells through inhibition of the mevalonate pathway within the cancer cells, but the reduction of circulating cholesterol levels through hepatic pathways is indeed considered important, especially for hormone-dependent cancers such as breast cancer ^24^. Cholesterol serves as a fundament building block for all steroid-based hormones, and disrupted estrogen synthesis might be beneficial for estrogen receptor (ER)-positive breast cancer. The cholesterol metabolite 27-hydroxycholesterol has been demonstrated to act as an ER-ligand that is able to promote progression of estrogen-dependent breast cancer ^25^. A recently conducted breast cancer trial demonstrated statins capability to effectively reduce circulating levels of 27-hydroxycholesterol ^20^, thus providing suggested mechanisms of the proposed systemic statin effects in obstructing breast cancer progression.

Some limitations of this study need to be acknowledged. This large-scaled nationwide study consisting of 20,559 breast cancer patients was limited to analyzing breast cancer related and overall mortalities, whereas considering endpoints including recurrences was not possible since data on breast cancer recurrences are not yet available on a national level in Sweden. In this study, results are based on a median follow-up time of 61.6 months, which allows for interpretations of statins’ effects regarding early clinical outcome of breast cancer, whereas later occurring mortality cannot currently be depicted in this study population. Moreover, there was no access to information regarding adjuvant treatment, which preferably should have been controlled for in the multivariate analyses. This drawback is somewhat balanced by the fact that all patients included in this study received their adjuvant treatment during a very limited period of time (2006?2008) and, importantly, no major changes in adjuvant treatment were introduced during these years (i.e. adjuvant trastuzumab was introduced in Sweden after this point). Additional data, which may have been of interest to control for, were co-medications such as aspirin. However, such medications can be dispensed over-the-counter and are not necessarily prescribed, which can lead to misclassification. Considering the associations between statin use and overweight ^26^ in addition to impaired breast cancer survival among overweight and obese individuals ^27^, it would have been preferred to control for body mass index as a confounder in the survival analyses. In parallel with the Danish study ^4^, statin use in this study was dominated by simvastatin, which may limit the generalizability to statins in general. In addition, the results from this study are based on data from a Swedish population and might not apply to other countries as indicated by the disparities between the previously published studies elsewhere in Europe and studies from the US. For example, adjuvant treatments for breast cancer may differ as a result of differences in healthcare systems, access to healthcare, and treatment guidelines. Furthermore, the Swedish national breast cancer screening program is well established and available for women between the ages 40 and 75 years. A recognized and highly assessed screening program is likely to be associated with early detection of breast cancer, resulting in lower stage disease at diagnosis, which will impact survival ^28^. Importantly, when interpreting the results of this study, a healthy user bias might have affected the results in cases in which statin use is associated with increased health awareness. This agrees with the likelihood of attending the screening program.

A couple of strengths of this study are noteworthy. Firstly, the considerable amount of breast cancer patients included in this study represents one of the largest studies of statins and breast cancer prognosis. Secondly, Swedish infrastructure allows for access to a high quality, national pharmaceutical database, cancer registry, and cause-of-death from which data were easily merged through the civil person number while prioritizing patients’ confidentiality as the procedures were handled by the Social Health and Welfare administration. Statins are prescribed drugs as described by the Swedish Prescribed Drug Register, and there should be a minimal risk of misclassification regarding statin use as over-the-counter administration is not possible. A possible exception would be if a patient had terminated statin use prior to July 1^st^, 2005 when the national register was initiated, however, we consider this unlikely since statins are usually prescribed as a lifelong treatment. Thirdly, all analyses were adjusted for a diagnosis of diabetes, which limits the possible survival bias that the diabetes might imply. Diabetic patients are commonly prescribed a statin in addition to their anti-diabetic medication, since diabetics have an elevated risk for cardiovascular diseases, which further highlights the need to consider diabetes in the analyses of breast cancer mortality. Finally, sensitivity analyses considering potential lag time bias used one-year lag-time based on calculations showing that after one year of statin use the effect reaches a plateau and is thus a reasonable lag-time estimate. Reassuringly, the effect estimates of statin use regarding both breast cancer specific survival and overall survival were strengthened in the sensitivity analyses taking lag time into consideration.

In conclusion, statin users, particularly simvastatin users, had a lower risk of breast cancer related deaths compared to non-users in this nationwide cohort of Swedish women with breast cancer diagnosed after the age of 40. Considering previous evidence from functional-, clinical- and epidemiological studies, this study adds evidence to the notion that statins possess beneficial effects against breast cancer progression along with its cardiovascular benefits. Above all, there is a need for confirmative results based on clinical trials before statin treatment can be recommended for patients newly diagnosed with breast cancer.

## Figure legends

**Supplemental Figure 1:** Lagged exposure and all-cause mortality. At every lag time marked on the x-axis the HR has been calculated. The figure suggests that the statin effect on mortality decreases with increasing lag.

**Supplemental Figure 2:** Truncated cumulative statin dose and all-cause mortality. Doses below each threshold have been set to zero. The figure indicates that the effect of statin treatment stabilizes after a cumulative dose of roughly 367 DDD.

